# Impact of Amazonian protected areas in preventing deforestation and carbon loss over four decades

**DOI:** 10.64898/2026.06.23.733928

**Authors:** Letícia Lopes Dias, Luiz Guilherme dos Santos Ribas, Bruno R. Ribeiro, Jonas Geldmann, Fabiana Prado, Neluce Soares, Paulo De Marco Júnior

## Abstract

Native vegetation protection is a key strategy for delivering both biodiversity and climate benefits, and protected areas have been widely adopted to keep tropical biomes standing. Yet deforestation is driven by interrelated environmental and social factors, and the effectiveness of protected areas varies considerably across space. Here, we evaluated the impact of 802 protected and conserved areas in the Brazilian Amazon on preventing vegetation loss and avoiding carbon emissions over the past 40 years using statistical matching to address the location bias of protection. We found that protected areas were effective throughout the study period, reducing the probability of deforestation per km^2^ by an average of 0.5 percentage points per year. While the Amazon biome lost 14% of its native vegetation between 1986 and 2024, protected areas prevented the deforestation of 290,436 km^2^, nine times their actual internal loss. They also stored 45,336 Mt of carbon in 2016 (61% of the Amazon stock) and prevented the emission of 7,300 Mt of CO_2_ by 2024. Deforestation inside the areas and remoteness reduced their impact, while areas that were initially more preserved were more effective. Area size and age had no influence over impact once we analyzed the amount of avoided deforestation per size and age. Impact also varied across Brazilian states, highlighting the role of regional context. All three protection categories (conservation units, indigenous lands, and quilombola territories) showed a positive mean impact, indicating that each, in aggregate, contributes to reducing deforestation. These findings provide robust evidence of the substantial role of Amazonian protected areas in habitat conservation and climate mitigation, while underscoring that this contribution remains undervalued. We advocate for strategically expanding protection to areas of greatest potential impact, and for securing adequate funding to ensure protected areas can fulfill that potential.

## Introduction

Preserving the Amazon plays a crucial role in mitigating the climate crisis, as the biome stores approximately 72,400 Mt of carbon, with 58% of this stored in protected areas, including indigenous lands (Walker et al., 2020). However, the region faces a series of threats that result in deforestation and degradation of the biome, affecting biodiversity, climate, and local communities (Lapola et al., 2023).

The main drivers of deforestation in the Amazon are the expansion of agriculture and livestock farming, mining, extreme climate events exacerbated by climate change, and political instability (Hänggli et al., 2023). Beyond direct causes, there are factors that indirectly influence deforestation, such as road construction and proximity to navigable rivers, which facilitate transportation and access to trading sites (De Souza & De Marco, 2014). Public policies and economic dynamics also indirectly affect deforestation, either by increasing the profitability of deforestation through rising commodity prices, or by raising its costs through enforcement, fines, and other penalties (Assunção et al., 2023).

Protected areas have a consistent effect in reducing deforestation due to their capacity to influence both the proximal and underlying causes of native vegetation loss (Hänggli et al., 2023). Understanding the variation in the capacity of these areas to generate concrete conservation outcomes can assist in designing more effective conservation strategies, since even with a sustained effect, they remain the conservation strategy with the smallest mean impact globally (Langhammer et al., 2024). Despite advances in evaluation, evidence on how conservation impacts vary across space and time, and how to downscale large-scale findings to local contexts, remains limited.

Most protected area impact evaluations rely on deforestation or other remote sensing data, as these have become increasingly available in recent decades (Ribas et al., 2020; Zhang et al., 2023). These datasets are particularly valuable for impact evaluations because they enable comparisons between areas inside and outside protected areas, the fundamental requirement for such assessments. They are also consistent and standardized over time, providing long time series suitable for robust evaluations (Souza et al., 2020). Yet, there is a lack of evidence on how to translate results from large-scales analysis to the site-level. Previous studies have measured the impact of Amazon protected areas on avoiding deforestation without addressing the drivers of impact (Alves-Pinto et al., 2022; Soares-Filho et al., 2010). On the other hand, global studies analyzing both impact and drivers generalize results to country level for feasibility (Shah et al., 2021; Yang et al., 2021).

By applying a counterfactual framework to assess the impact of Amazonian protected areas on preventing deforestation over the past 40 years, we estimated site-level impact and examined how it relates to protected area characteristics and spatial context. We also quantified avoided carbon emissions from 2016 to 2024, constrained by carbon data availability. We combined deforestation and carbon data to estimate protected areas’ impact on one of the major causes of biodiversity decline and the climate crisis, advancing on identifying spatial and temporal patterns of effectiveness. Our findings offer actionable recommendations to strengthen management and maximize the contribution of protected areas to both biodiversity conservation and climate action.

## Methods

### Study area

This study analyzes conservation units, indigenous lands, and quilombola territories within the Brazilian portion of the Amazon biome, categories recognized as protected areas under the National Strategic Plan for Protected Areas (National Strategic Plan for Protected Areas, 2006). The Amazon has more than half of its extent covered by protected and conserved areas, making it the Brazilian biome with the greatest protection coverage (Alves-Pinto et al., 2022). Here, we use georeferenced data available from the National Registry of Conservation Units (CNUC; MMA, 2025), the database of the National Foundation for Indigenous Peoples (FUNAI, 2024), and the National Institute for Colonization and Agrarian Reform (INCRA, 2023). We excluded from the analysis indigenous lands currently under study (i.e. areas under analysis to become legally protected) in FUNAI’s database, and quilombola territories with annulled titles in INCRA’s database, as these are not legally formalized and therefore lack the appropriate management structure to be officially considered protected. Based on these criteria and the availability of valid spatial data, our sample comprised 802 protected areas, being 364 conservation units (92% of those registered in the CNUC within the biome), 317 indigenous lands (95% of those registered with FUNAI), and 121 quilombola territories (88% of those registered with Incra; Figure S1; Table S1).

Data on the extent of each protected area, originally in vector format, were rasterized at a 1 x 1 km resolution using the *terra* package in R (Hijmans, 2020). A cell was considered “protected” when at least 75% of its area was covered by a protected area polygon. Conversely, cells from 0 to 25% of protection coverage were considered non-protected, and so, available to be selected as control units in the matching process. For the analysis, we separated the areas by year (1985 to 2024), grouping areas with a creation year equal to or earlier than the analysis year into a single raster file. For example, in the 2012 protected areas raster, cells classified as protected refer to those in which the areas had been delineated in 2012 or earlier.

We began our analysis by preprocessing data on land use and land cover, protected areas, and matching covariates, converting vector and raster inputs into standardized raster data at a 1 × 1 km resolution (Figure 1). The rasters were then transformed into data frames for each analysis year, in which each row corresponded to a cell and recorded whether the cell was protected, whether it was deforested in that year, and the values of each covariate. These datasets served as inputs for the matching procedure, which produced balance estimates and treatment-group assignments for each cell. The analysis then followed two complementary directions. First, we estimated annual avoided deforestation at the biome level. Second, we linked cell-level results back to their corresponding protected areas to estimate avoided deforestation for each site. Building on these site-level estimates, we incorporated carbon data to quantify yearly avoided emissions per protected area, and combined them with site attributes to analyze the factors associated with the estimated impact.

**Figure 1.**
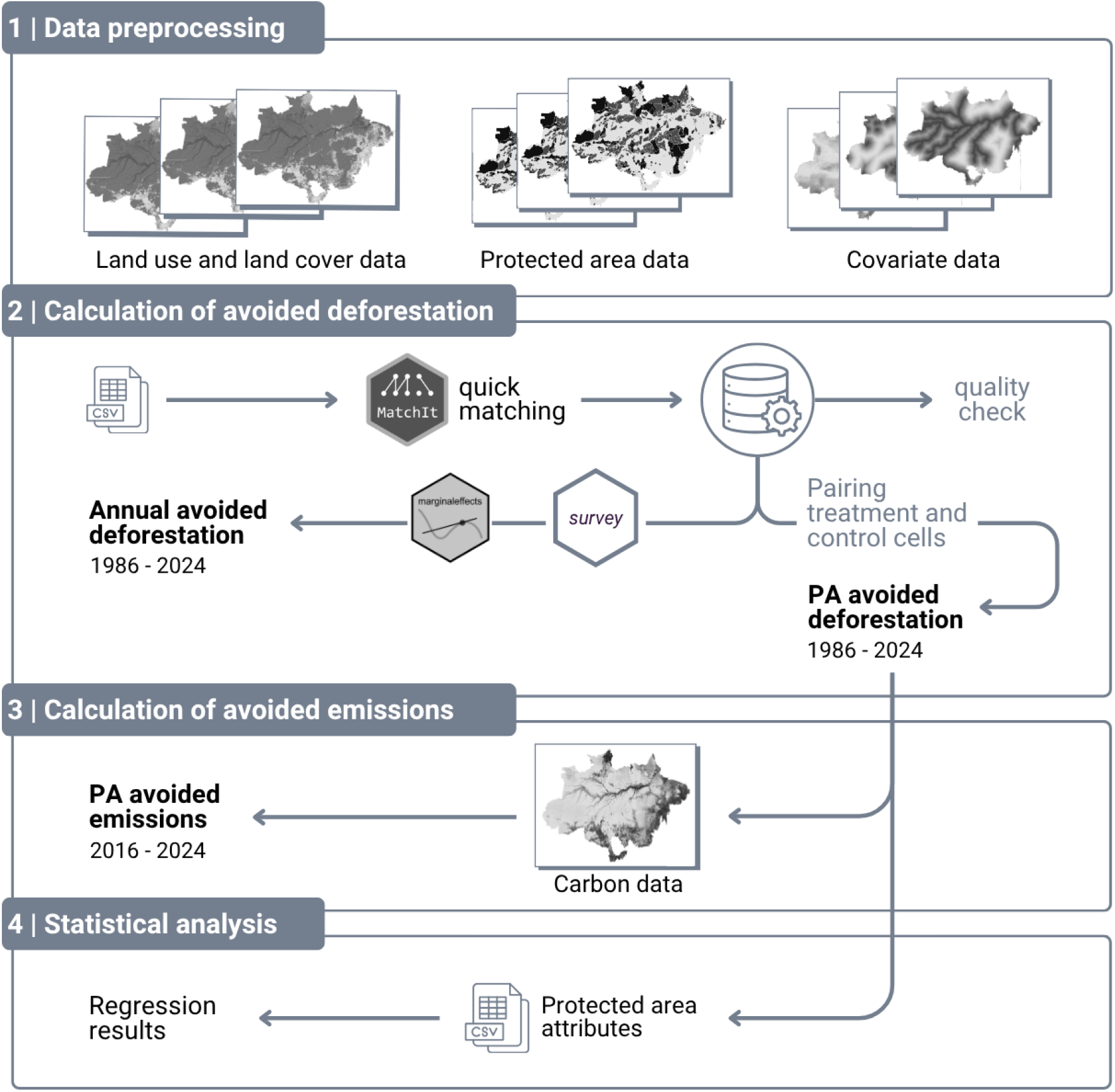
Methods flow chart. PA = protected area.

### Land use and land cover data

We used land use and land cover data from the 10^th^ collection of the MapBiomas Project (MapBiomas, 2025; Souza et al., 2020), covering the period from 1985 to 2024. Data was aggregated from original 30 x 30 m resolution to 1 x 1 km and downloaded through the Google Earth Engine platform, on August 18, 2025. When aggregating, we used the mode function, so the cell kept the class of the most frequent land use and lad cover class. Then, we used R software (R Core Team, 2024) to calculate deforestation that occurred from 1986 to 2024 by reclassifying raster data into: native vegetation (such as forest formation, savanna formation, or mangrove) or non-native vegetation (such as pasture, agriculture, urban infrastructure, etc.). Cells classified as native vegetation in the previous year and as non-native vegetation in the following year were considered “deforested”.

### Calculation of avoided deforestation from 1986 to 2024

To assess the impact of protected areas in avoiding deforestation, we employed a matching method to compare protected cells to similar unprotected cells. The analysis involves comparing the outcomes observed in these areas with the hypothetical scenario of no protection (that is, the counterfactual scenario), constructed using the statistical matching technique based on covariates associated with deforestation and territorial protection (Ribas et al., 2021). Matching reduces the location bias of existing protected areas on their outcome in containing native vegetation loss (Ribas et al., 2021).

Deforestation is driven by proximal causes (such as agriculture, infrastructure expansion, and mining) and underlying causes, including economic, social, and institutional factors that indirectly fuel the former (Hänggli et al., 2023). Understanding these pathways allows us to develop a theory of change (Figure 2) in which the impact of protected areas on deforestation can be assessed by controlling for confounders through statistical matching (Ribas et al., 2021).

Drawing from the literature, we selected covariates related to land use change and to the delineation of an area for protection (Figure S2; Table 1). Covariates were standardized to raster format at a 1 x 1 km resolution.

**Table 1.**
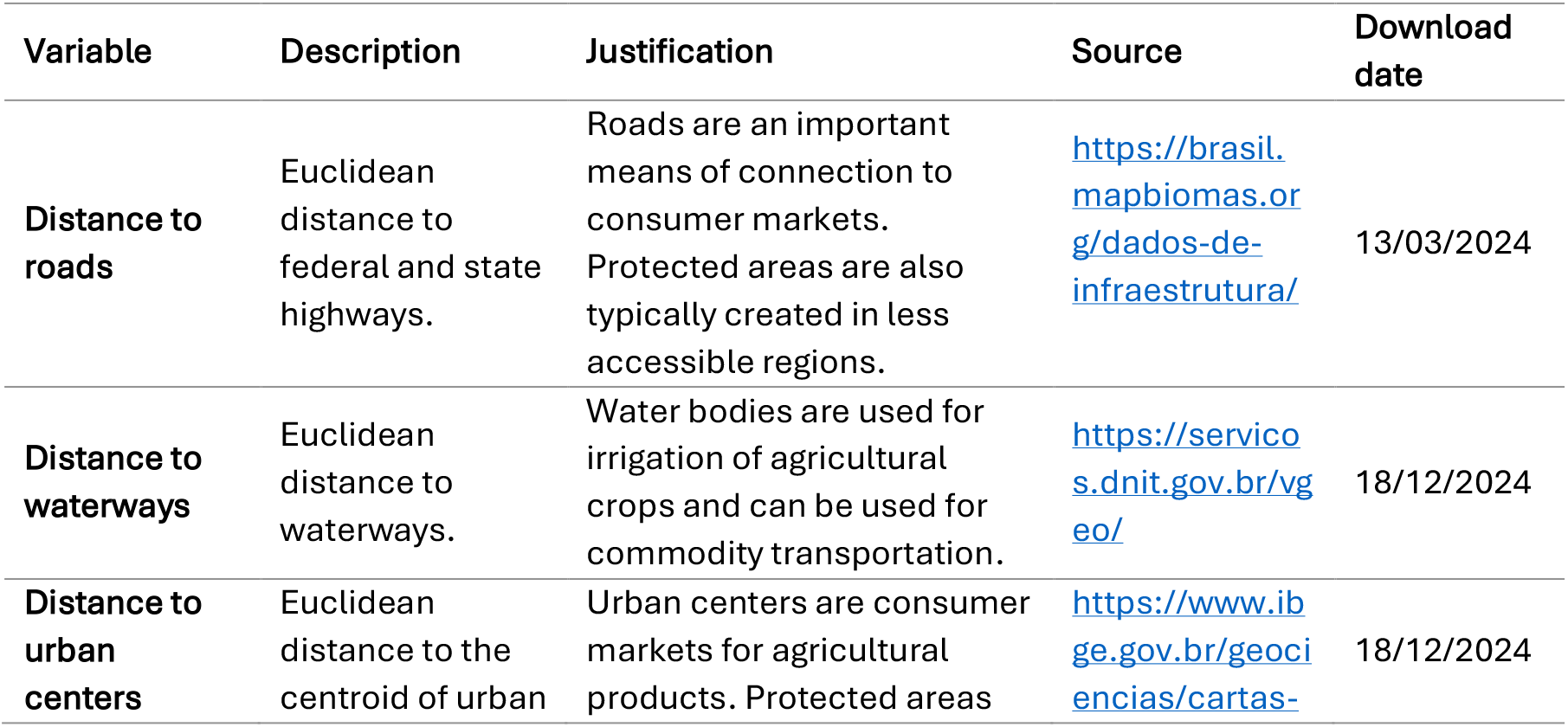
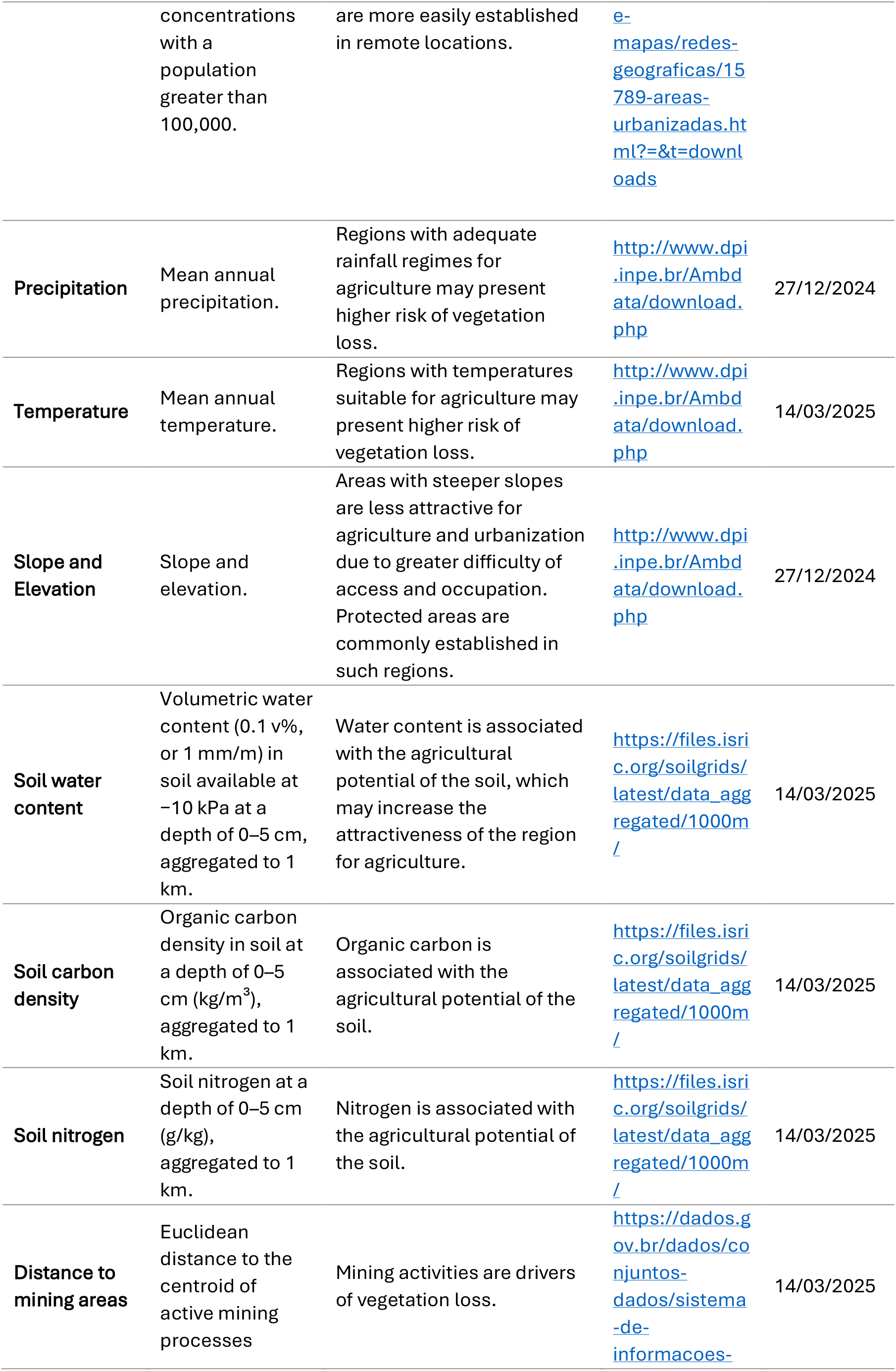
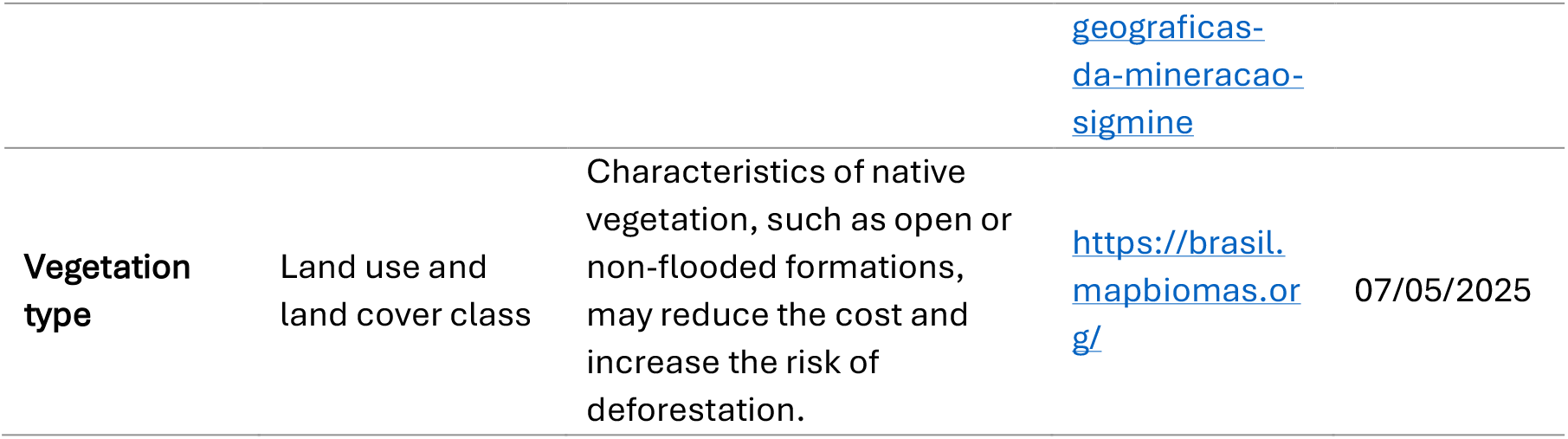
Variables used for statistical matching between protected and unprotected cells.

For matching, we used the *MatchIt* package in R (Ho et al., 2011) and followed the protocol proposed by Ribas et al. (2021). We applied the generalized full matching approach, which optimizes the matching process and is well-suited to large datasets (Ho et al., 2011). Given the scale of our analysis (encompassing the Amazon biome across a 40-year period) this method was particularly suitable, as it produces optimal matched solutions without the prohibitive computational demands associated with other matching algorithms (Sävje et al., 2021). We then compared the mean difference on covariates between protected and unprotected groups to assess matching quality. Matching is considered successful when there is a significant reduction in the mean difference between covariate values for treated (protected) and control (unprotected) cells (Sekhon, 2011).

We structured the dataset resulting from matching as a weighted survey design using the *survey* package in R (Lumley et al., 2024), and calculated the annual treatment effect using survey-weighted generalized linear models. This approach adequately incorporates matching weights, given that full matching allows replacement, meaning that the same control unit can be used as a comparison for multiple treated units. We then estimated the average treatment effect on the treated (ATT) using the avg_comparisons() function from the *marginaleffects* package (Arel-Bundock et al., 2024). Finally, the total area of avoided deforestation per year was calculated by multiplying the regression coefficient by the sum of protected cells.

In addition to the avoided deforestation results for each cell, we calculated outcomes at the protected area level. To do so, we defined the control group from the dataset resulting from matching, which includes subclass values and the probability of a cell being protected. Within each subclass, we selected the unprotected cells with the highest probability, replicating a cell when the subclass contained more treatment than control cells. We then matched the selected control cells with their corresponding protected cell belonging to the same subclass. The result was a table of paired protected and unprotected cells in equal numbers. Avoided deforestation was calculated as the difference between the observed deforestation in the protected cell and in the control (unprotected) cell. Finally, we identified which protected area each cell corresponded to and summed the avoided deforestation across cells within each protected area, obtaining the avoided deforestation per protected area and per year since its establishment until 2024. For area-level analysis, we kept cells belonging to more than one exclusive site, due to some geographical overlaps; while for biome estimations we removed overlaps.

The generalized full matching method allowed adequate matching of treatment cells (i.e., protected) to control cells, keeping most standardized mean differences for covariates below 0.1 in analyses up to 2005 (Figure S3). From 2006 onward, eight covariates exceeded the 0.1 threshold; three remained below 0.25. Despite these variations, matching was effective in reducing the difference between treated and control units across all years analyzed. Following the matching procedure, we assessed the geographic distance between paired protected and control cells. Distances ranged from 0 to 30 km, with a plateau in the frequency distribution between 1 and 18 km, indicating a homogeneous concentration of control cells within that range (Figure S4).

### Calculation of avoided carbon emissions from 2016 to 2024

To quantify the CO_2_ emissions avoided by protected areas in the Amazon from maintaining forest cover, we used the biomass map produced by Ometto et al. (2023), downloaded on April 16, 2025, through a project of the National Institute for Space Research (INPE). This map is based on LiDAR (Light Detection and Ranging) data, forest inventories from partner institutions, and satellite data from 2016 (Ometto et al., 2023). Additionally, biomass in other compartments, such as roots, deadwood, and litter, was estimated by aggregating information from the phytophysiognomies of the historical natural vegetation map of the Amazon and applying above-ground biomass expansion factors (Ometto et al., 2023). This map has been used as the official estimate by the Brazilian government.

We estimated avoided emissions by calculating the biomass in each protected cell and multiplying it by the avoided deforestation value from 2016 onward. We then obtained values per protected area by pairing protected cells with control cells from the same subclass with the highest probability of being protected. We considered the maximum potential carbon loss that could be emitted into the atmosphere, assumed to occur entirely at the moment of deforestation, corresponding to 100% of the initial total carbon stock (Achard et al., 2014; Song et al., 2015). Finally, avoided CO_2_ emissions were estimated by multiplying total avoided carbon loss by 3.67, the molecular weight ratio of CO_2_ to carbon (CO_2_/C).

### Statistical analysis

To investigate which factors were associated with protected area impact on curbing deforestation, we fitted a linear regression model with avoided deforestation rate as the dependent variable. The deforestation avoidance rate (*def*_*rate*) for each protected area was calculated as:

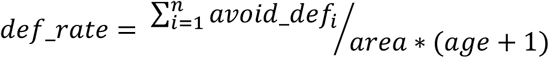

where 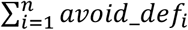 is the cumulative avoided deforestation (in km^2^) summed across all years *i* within the study period, estimated as the difference between the deforestation in unprotected control cells and the deforestation within the protected area boundary; *area* is the total area of the protected area (km^2^); and (age + 1) is the number of years since the protected area was established, incremented by one to prevent division by zero for newly created units. The resulting index expresses the annual avoided deforestation per unit area, standardizing for both the size and the age of each protected area to allow comparisons across sites.

As explanatory variables, we included the observed deforestation rate over the same period; area size (in km^2^), age (in years), and type (i.e., indigenous land, conservation unit, or quilombola territory); the proportion of native vegetation at the time of establishment; the Brazilian state with which the area most overlaps; and the mean Euclidean distance to urban centers. Protected areas represented by fewer than three cells were excluded from this analysis, as estimates based on one or two cells are unreliable and likely to introduce substantial bias in the impact rate calculation. As a result, the final model sample comprised 725 protected areas while maintaining 99% of the total coverage of protection within the Brazilian Amazon.

As for avoided deforestation rate, we estimated the observed deforestation rate by dividing the total deforestation that occurred within each area by its surface area and age. The proportion of native vegetation at establishment was calculated using land use and land cover data for the year each protected area was created. Native and non-native vegetation were classified from 1 km-resolution rasters, and the proportion of native vegetation was obtained by dividing the number of native cells by the total number of cells within each area.

After fitting the model, we assessed residual normality and multicollinearity among predictors. Residual plots indicated heteroscedasticity, meaning that the variance of residuals was not constant across fitted values. To obtain reliable inference under these conditions, we re-estimated standard errors and p-values using heteroscedasticity-consistent (HC1) standard errors with state-level clustering, applied via the tab_model() function from the *sjPlot* package (Lüdecke, 2018). This approach adjusts the covariance matrix of the coefficients to account for non-constant residual variance and potential within-state correlation among observations, without requiring model refitting.

Additionally, we assessed the relationship between avoided deforestation and avoided carbon emissions between 2016 and 2024 across protected areas applying a Pearson’s correlation test. All analyses were conducted in R (R Core Team, 2024).

## Results

From 1985 to 2024, protected areas prevented the deforestation of 290,436 km^2^, an area equivalent to approximately 80% of Germany’s territory. Over the same period, these areas lost 32,067 km^2^ of native vegetation internally, meaning that, in the absence of protection, deforestation within these boundaries would have been approximately nine times greater. Meanwhile, the entire Amazon lost 735,815 km^2^ of native vegetation, equivalent to 14% of the total area (Figure S5). This loss was concentrated in the southern and eastern portions of the biome, “the deforestation arc” (Velasco Gomez et al., 2015), and mostly driven by the conversion pastures.

Indigenous lands, covering 1,058,074 km^2^, avoided the conversion of 173,363 km^2^; conservation units, covering 1,178,649 km^2^, avoided 136,116 km^2^; and quilombola territories, covering 17,577 km^2^, avoided 3,342 km^2^.

Protected areas were effective in containing deforestation (Figure 4a) in all years (Figure 4b). The average effect of protection, however, varied over time: it increased until 2003, declined in the following years, and stabilized between 2010 and 2018; the effect then rose again between 2019 and 2022 (Figure 4c). The average treatment effect ranged from −0.87% (in 2003) to −0.05% (in 1986), reducing on average the probability of deforestation per km^2^ by 0.5% per year.

**Figure 3.**
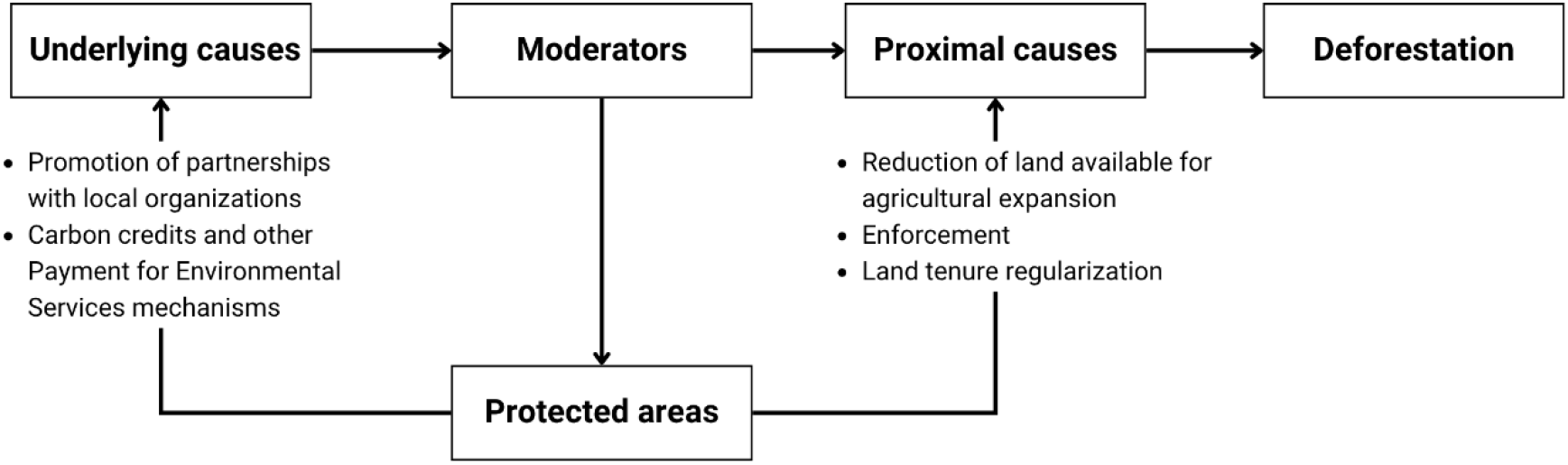
Relationship between underlying causes, moderators, and proximal causes of deforestation, and the role of protected areas as conservation instruments within this dynamic. Moderators are the mechanisms through which underlying causes affect proximal causes and public deforestation control policies. The topics highlight examples of how protected areas can act upon the causes of deforestation. Adapted from: Hänggli et al. (2023).

**Figure 4.**
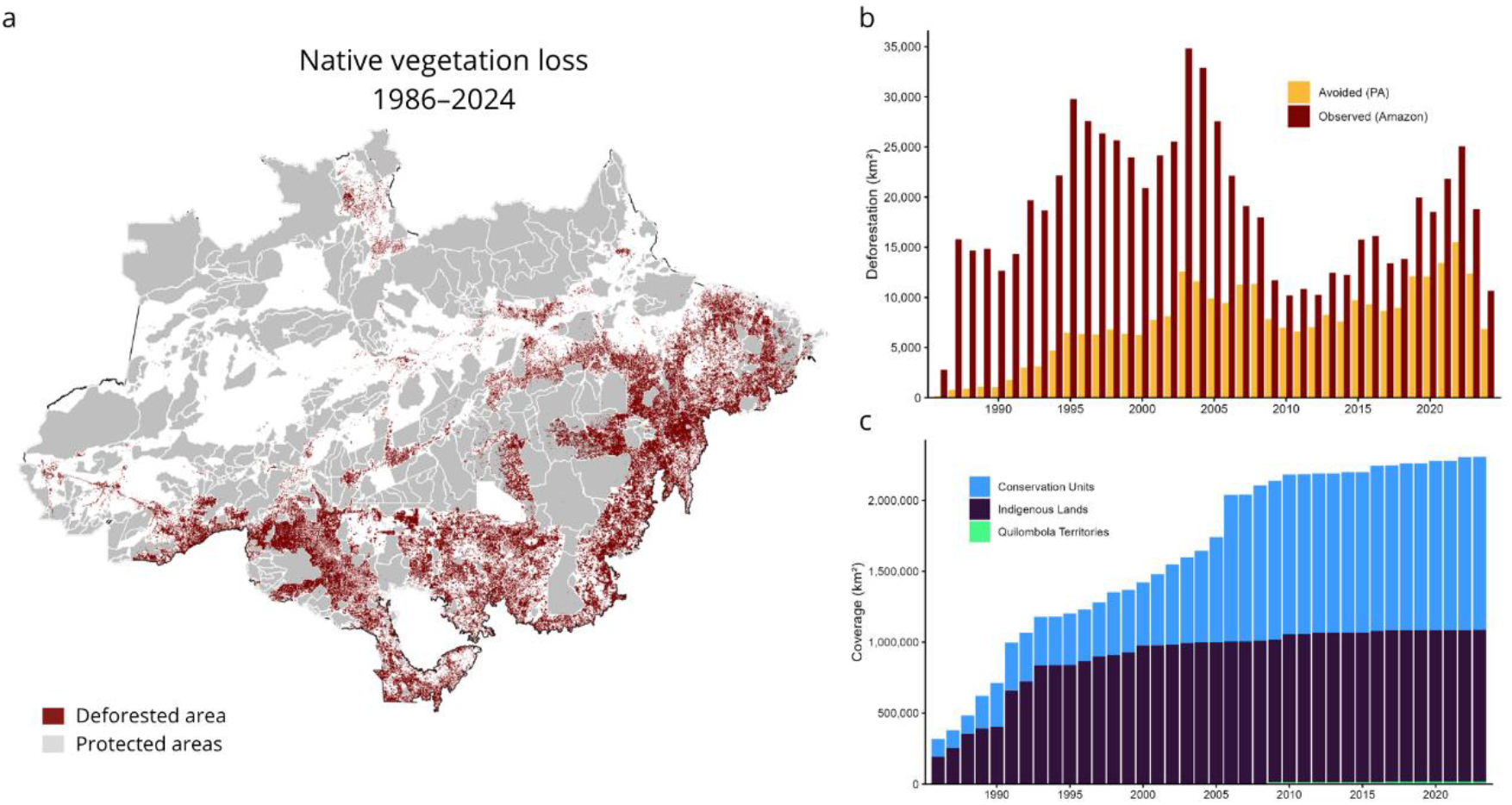
Native vegetation loss and estimated effect of protected areas in the Amazon biome. a) Spatial distribution of cumulative native vegetation loss between 1986 and 2024, highlighting protected area boundaries as barriers to deforestation. b) Total annual deforestation (red bars) and avoided deforestation attributed to protected areas (orange bars), in km^2^, over the analyzed period. c) Annual coverage of protection, according to protected area type.

When examining the impact in terms of avoided deforestation per unit area and per age, of the 802 areas analyzed, 73% had a positive impact; 9% had a negative impact (meaning they exhibited higher deforestation than their matched control areas) and 19% had no detectable impact (Figure 5a), either because no deforestation occurred in the control areas or because it occurred at the same rate as in the protected area (Figure 5b; Table S2).

**Figure 5.**
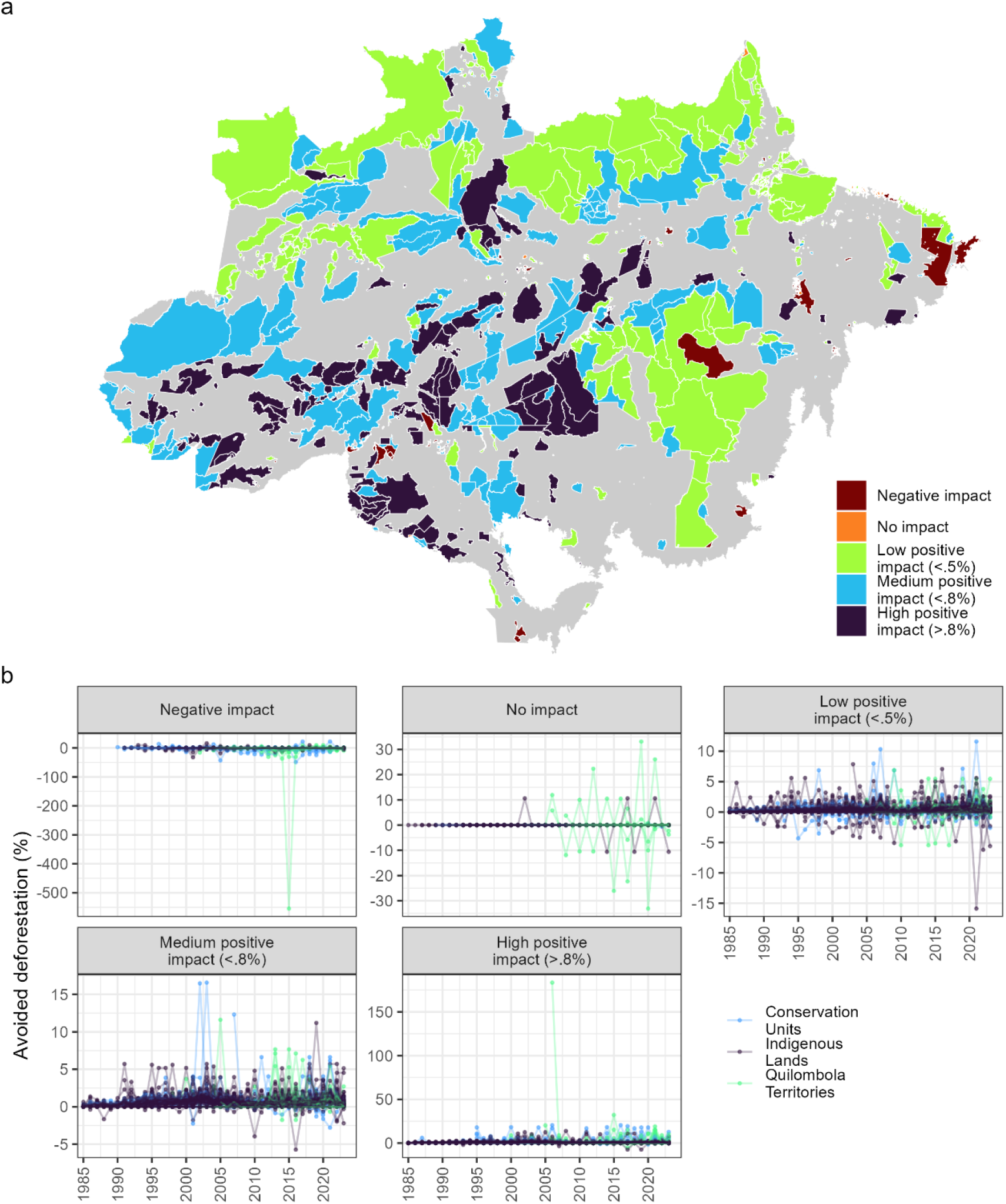
a) Avoided deforestation rate by protected areas from 1986 to 2024. b) Temporal variation in avoided deforestation rate per protected area, by type and magnitude of impact rate. The impact rate was calculated by dividing avoided deforestation by the extent of the protected area (km^2^), divided by the age of the area. Magnitude was classified as: negative impact = rates < 0; low positive impact = rates < lower quartile (0.5%); medium impact = rates between the lower quartile (0.5%) and the median (0.8%); and high positive impact = rates > median.

Protected areas stored a total of 45,336 Mt of carbon in 2016 (Figure 6a), which represents 61% of the total Amazon stock. From that year through 2024, they prevented the loss of 93,856 km^2^ of native vegetation, resulting in the mitigation of 1,997 MtC in emissions, assuming 100% conversion of the original biomass (Figure 6b). This represents 7,300 Mt of CO_2_ of avoided emissions. Pearson’s test indicated a strong, positive correlation between avoided deforestation and avoided carbon emissions over the same period (r = 0.953; 95% CI [0.946, 0.959]; p < 0.001).

**Figure 6.**
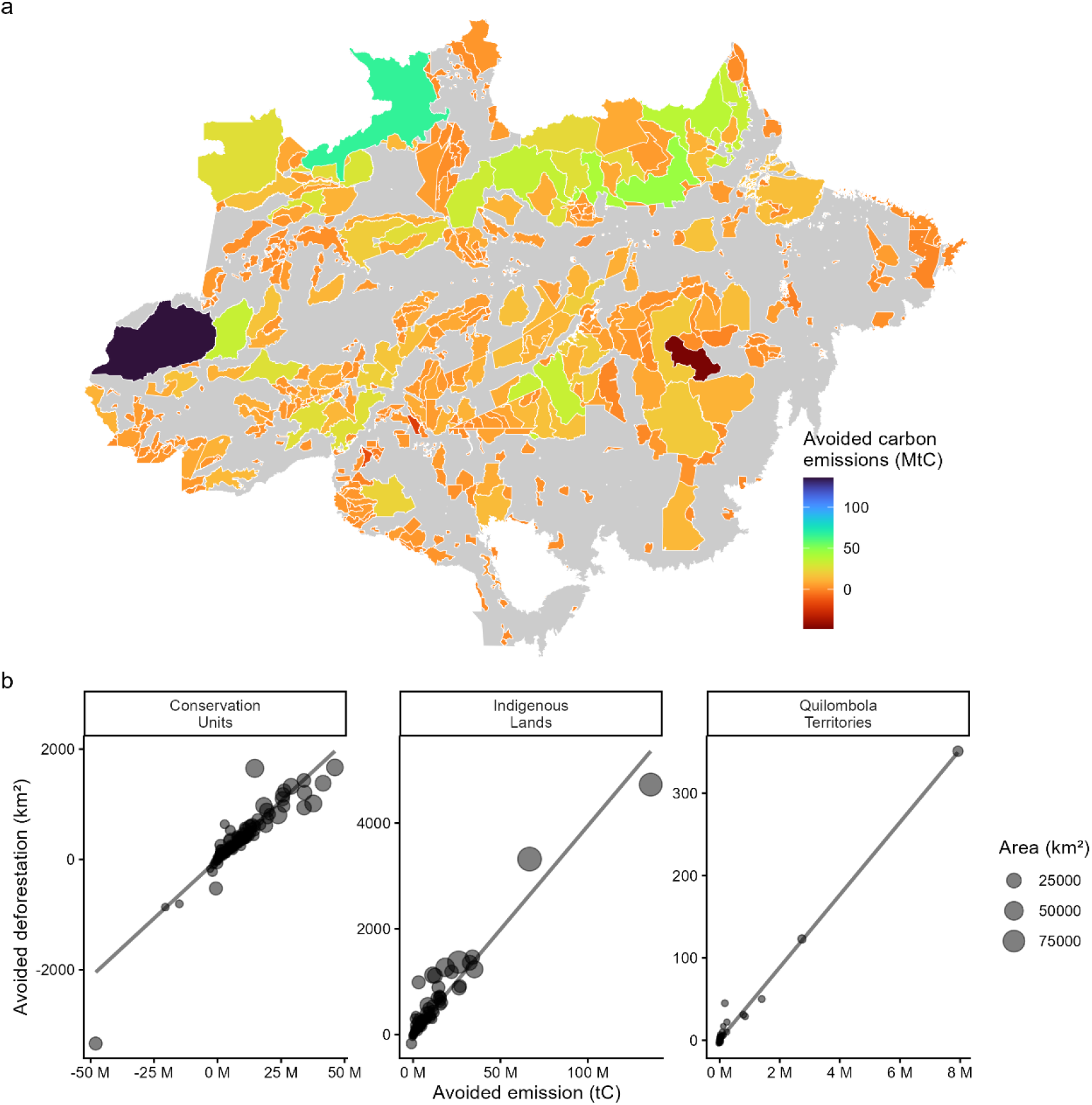
a) Total carbon emissions avoided by protected areas from 2016 to 2024; b) Relationship between avoided deforestation (km^2^) and avoided carbon emissions (tC) for protected areas between 2016 and 2024. Each point represents one protected area, with point size proportional to area. Lines represent fitted linear regressions.

**Figure 7.**
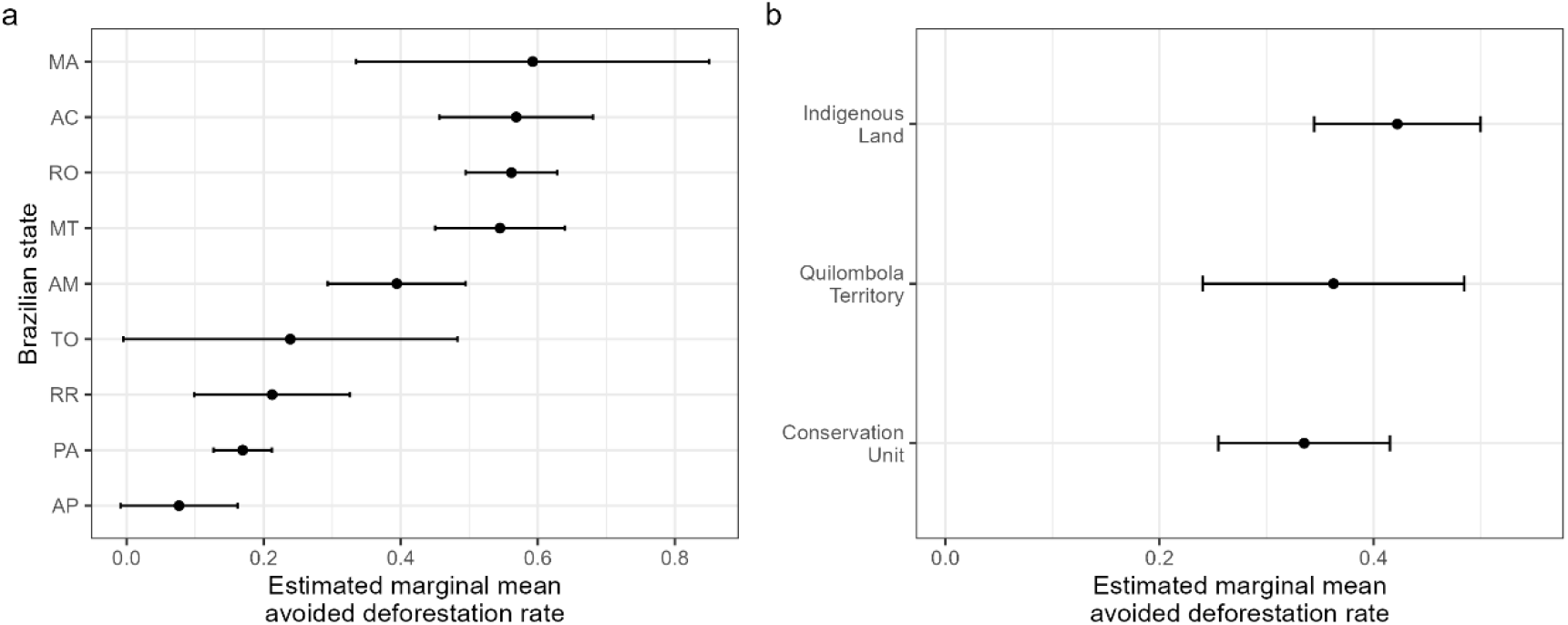
Estimated marginal mean avoided deforestation rate by (a) Brazilian state and (b) protected area type, derived from a linear regression model. Points represent estimated marginal means, and horizontal bars indicate 95% confidence intervals. State abbreviations refer to the Brazilian states of Acre (AC), Amapá (AP), Amazonas (AM), Maranhão (MA), Mato Grosso (MT), Pará (PA), Rondônia (RO), Roraima (RR), and Tocantins (TO).

Protected area impact, measured as the avoided deforestation rate, was negatively associated with the observed internal deforestation rate (β = −0.718, p < 0.001) and distance to urban centers (β = −0.0004, p < 0.001), and positively associated with the initial proportion of native vegetation (β = 0.006, p = 0.010; Table S3). These suggest that areas facing higher actual deforestation tend to have lower impact, as do areas located farther from urban centers, where deforestation threat is inherently lower, and those established in already-degraded landscapes.

Impact rate also varied significantly across Brazilian states (non-overlapping confidence intervals in Figure 6a), suggesting that regional context plays a role in shaping protected area effectiveness. In contrast, no significant differences were found among protected area types (Figure 6b); nevertheless, all three types showed a positive mean impact, indicating that legal protection, regardless of its form, consistently contributes to reducing deforestation.

## Discussion

The impact of protected areas was consistently positive throughout the 40 years of analysis, reducing the probability of deforestation and carbon emissions relative to unprotected areas. We were able to conduct an analysis over a broader temporal scale than previous studies (Alves-Pinto et al., 2022; Gonçalves-Souza et al., 2021; Soares-Filho et al., 2010) and advance on site-level evaluation. We also found a strong overall effect: protected areas prevented approximately nine times the amount of deforestation they experienced internally. This stands in contrast to a global estimate in which protected areas prevented only 30% of the deforestation that occurred inside those areas (Yang et al., 2021), and may reflect the particularly high effectiveness of Amazonian protected areas documented at both national (Gonçalves-Souza et al., 2021) and global scales (Yang et al., 2021).

Our results advance the discussion on the role of protected areas in carbon storage by quantifying the carbon stock that would have been lost in their absence (Soares-Filho et al., 2010; Walker et al., 2020). Land use change is the major driver of carbon emissions in Brazil, and without the protection of those sites we estimated that national emissions between 2016 and 2024 would have been almost 50% greater, considering that actual net emissions were around 15.7 thousand Mt CO_2_ (Climate Observatory, 2026). In the same period, Brazil received approximately US$ 651 million through the Amazon Fund for avoiding the emission of 149 Mt CO_2_, roughly US$ 4 per ton (MMA, 2026). Our estimates suggest that the avoided emissions attributable to these protected areas may be nearly 49 times greater than the volume compensated through the Amazon Fund over the same period. This underscores the potential of robust impact assessments to ensure that the full climate contribution of protected areas is properly recognized and rewarded.

Initially, the amount of avoided deforestation increased along with protection coverage. However, following the overall reduction in deforestation across the biome, their effect stabilized, rising again as deforestation pressure intensified. An important milestone to increase protection coverage was the elaboration of the Action Plan for the Prevention and Control of Deforestation in the Legal Amazon (PPCDAm), launched in 2004 by the federal government which led to the doubling the extent of protected areas in the Amazon over a decade (Assunção & Gandour, 2018). Furthermore, these areas were strategically allocated in the region of greatest pressure, known as the “deforestation arc,” to form a shield against devastation (Assunção & Gandour, 2018).

Larger protected areas showed greater impact in terms of the amount of deforestation avoided and, consequently, avoided carbon emissions. The Vale do Javari (in Amazonas) and Yanomami (spanning Roraima and Amazonas) indigenous lands rank among the largest protected areas and showed the greatest impacts against deforestation, protecting 28,650 km^2^ from conversion. However, when measuring impact per size and year, larger areas did not show a higher effect. To contribute to the longstanding debate over the best strategy for establishing protected areas — multiple small areas or fewer but larger ones (Armsworth et al., 2018) — we add that while large areas stand out in absolute value, smaller areas remain effective and collectively contribute to the mitigation of carbon emissions. Previous studies also highlight that the protection of large fragments offers greater cost-effectiveness at reducing fragmentation and may also preserve habitat remnants capable of sustaining viable populations (Cho et al., 2019; Pimm et al., 2018). Thus, larger areas can combine ecological and climate benefits, since effectiveness at mitigating emissions and curbing deforestation is strongly related. Smaller sites, on the other hand, offer greater cost-effectiveness when the aim is to protect a larger number of species, and numerous small protected areas are located in biodiversity hotspots (Cho et al., 2019; Pimm et al., 2018). We cannot disregard the contribution of smaller sites, but our evidence supports that prioritizing larger remnants may offer greater combined benefits for both climate and biodiversity.

Regardless of their size, protected areas established in remote regions face inherently lower deforestation threats, and therefore have less potential deforestation to prevent (Yang et al., 2021). Yet these regions are precisely the ones traditionally favored, as they represent a lower cost of establishment (Vieira et al., 2019). Along with the lack of resources, this location bias may explain why this conservation strategy remains proportionally less effective than other measures, such as invasive species control and habitat restoration (Langhammer et al., 2024). As protected area impact is commonly measured through native vegetation indicators, as we did here, evidence of whether such areas deliver broader conservation outcomes remains limited.

Areas experiencing higher internal deforestation were also less effective, suggesting that when threat levels exceed a protected area’s capacity to respond, effectiveness declines. This points to the role of local management capacity (Powlen et al., 2021; Pulido-Chadid et al., 2025), which we were unable to test due to the lack of comprehensive data. These findings suggest that while some level of pressure is necessary for a protected area to demonstrate impact on curbing deforestation, excessive pressure, in the absence of adequate management, may undermine it. Thus, the establishment of new protected areas should be guided not only by the potential to generate conservation impact, but also by a commitment to providing the management conditions necessary for that potential to be fully realized. Additionally, some levels of deforestation are legally allowed and expected in certain categories of protected areas, except for strict protection conservation units. For quilombola territories and indigenous lands, forest conservation is largely a consequence of sustainable livelihood practices rather than the primary goal of their establishment (Zhang et al., 2023).

Regarding protection categories, no differences in avoided deforestation rate were found when controlling for size, age, and location. This suggests that indigenous lands, conservation units and quilombola territories are equally effective, contrasting with previous studies reporting differences in effectiveness between protection types (Alves-Pinto et al., 2022; Nolte et al., 2013). In Australia, a comparison between different levels of protection demonstrated that multiple-use areas were as effective as more restrictive ones in protecting forest cover (Stoudmann et al., 2025). Meanwhile, a global analysis found that more restrictive areas tend to be more effective for forest conservation (Yang et al., 2021). Despite extensive evidence of the importance of territories managed by traditional peoples and communities (Nolte et al., 2013; Ribas & Galetti, 2025; Walker et al., 2020), understanding the effect of governance, in a broader sense, on conservation outcomes remains a challenge. Zhang et al. (2023) reviewed the literature on this matter and, when relying on quantitative assessments, found that there was no governance category consistently better at reducing forest loss than others. Differences reported in previous studies may stem from direct comparisons between governance types without accounting for confounding factors such as size, age, and the administrative and political context in which protected areas are embedded.

We recognize limitations that should be considered when interpreting our results. First, native vegetation cover is an indirect measure of conservation outcome, and future studies should incorporate comprehensive biodiversity data, once they are made available, to fully capture the ecological impact of protected areas (Ferraro & Hanauer, 2015). Second, the absence of management data prevented us from isolating the effect of specific management attributes on effectiveness; an important next step, but one that must be pursued in integration with the structural factors identified here, to build a complete picture rather than capturing isolated pieces. Finally, while matching reduces location bias, residual confounding from unmeasured variables cannot be entirely ruled out.

Our results add to the growing body of evidence on the importance of protected areas for safeguarding natural ecosystems and mitigating the climate crisis (Geldmann et al., 2019; Ribas & Galetti, 2025; Shah et al., 2021; Shi et al., 2020), contributing to the understanding of the temporal and geographical patterns of their effectiveness. By combining robust analytical tools with available data, we provide meaningful evidence to enhance protected area effectiveness through two complementary strategies: strengthening currently established areas and expanding protection coverage. For currently established areas, urgent action is needed where impact is negative or declining: since higher internal deforestation is associated with lower effectiveness, reducing those losses must be the immediate priority. For new protected areas, more strategic planning is needed: designations should be located to safeguard remaining native vegetation in the most threatened regions, with sufficient resources allocated for effective management from the outset.

## Acknowledgements

This work was supported by the Gordon and Betty Moore Foundation through the LIRA (Amazon Region Integrated Legacy Initiative) project, led by IPÊ – Institute for Ecological Research, and by the Coordenação de Aperfeiçoamento de Pessoal de Nível Superior – Brasil (CAPES).

## Notes

https://zenodo.org/records/20507618

### Competing Interest Statement

The authors have declared no competing interest.

