## Supplemental Information for "Impact of Amazonian protected areas in preventing deforestation and carbon loss over four decades"

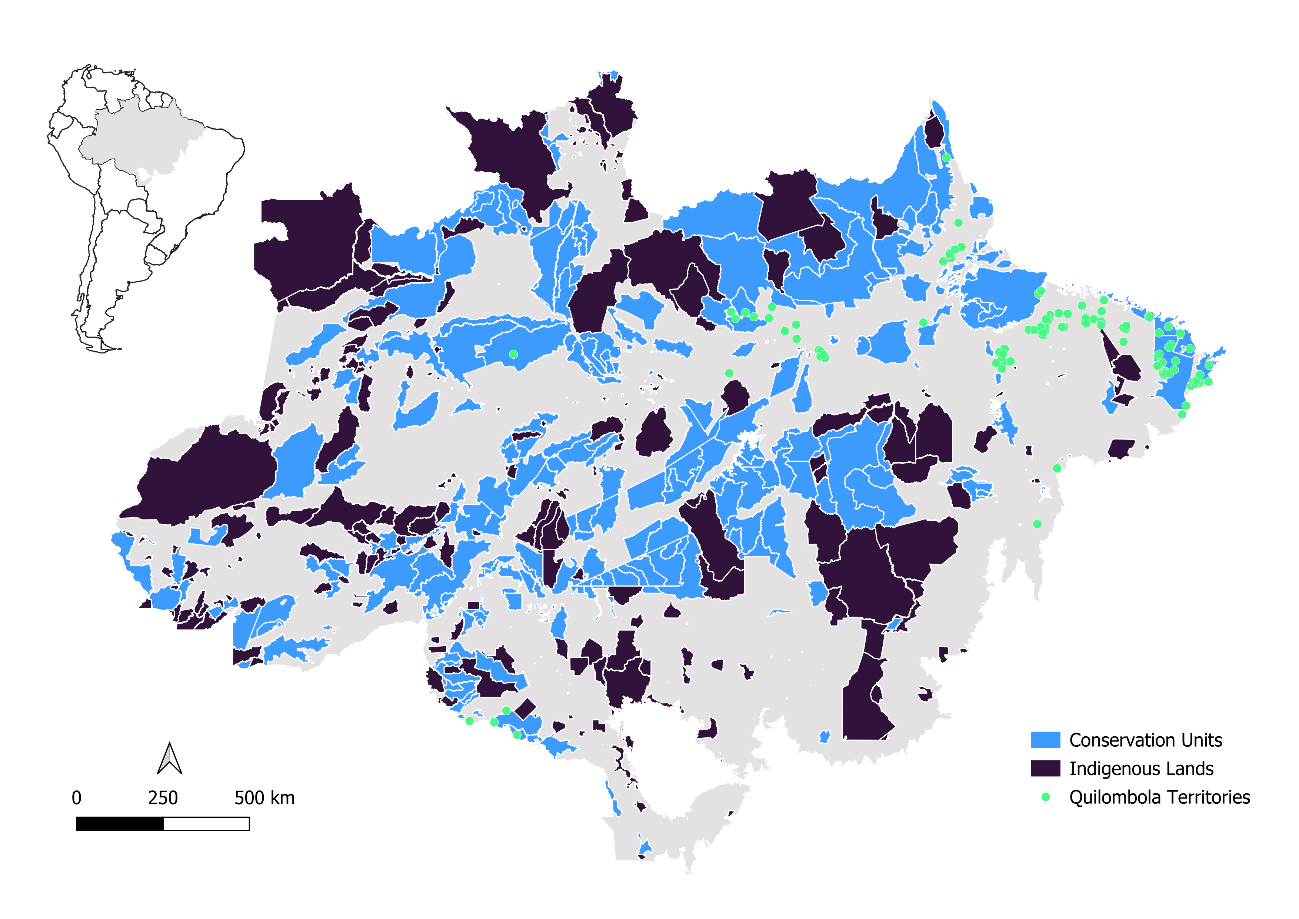

Figure S1. Spatial distribution of the protected areas analyzed in this study, located within the Brazilian Amazon biome.

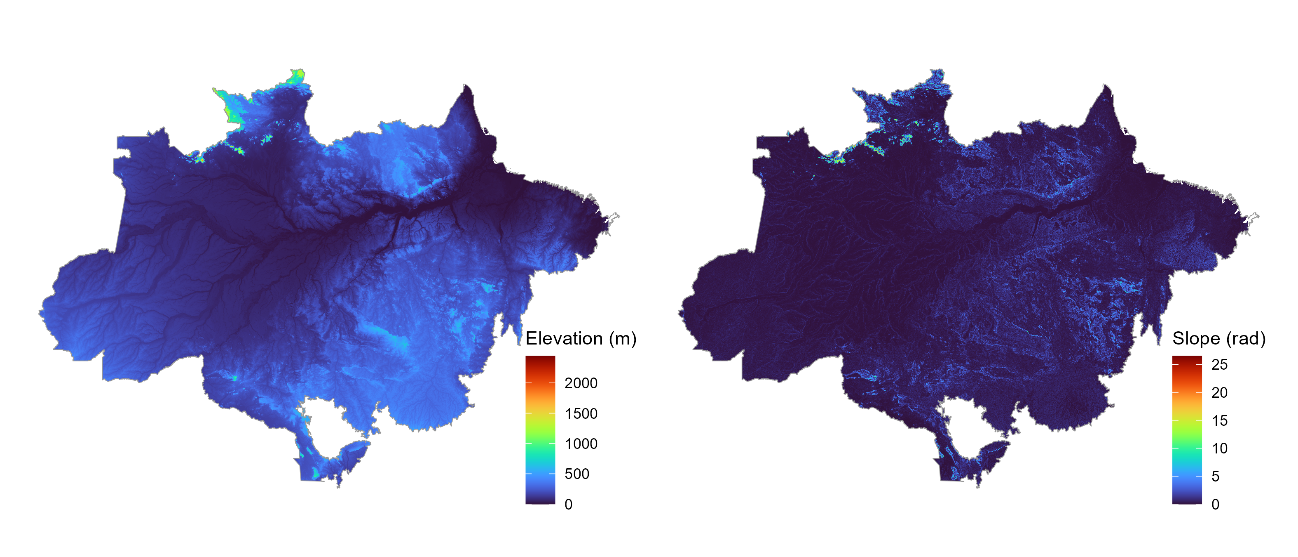

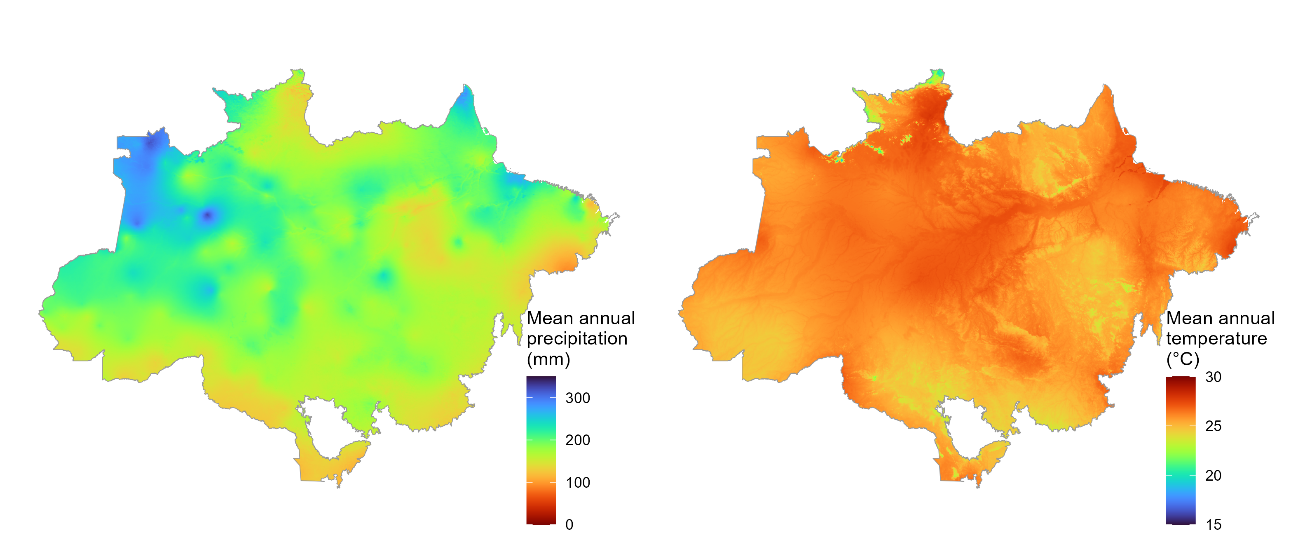

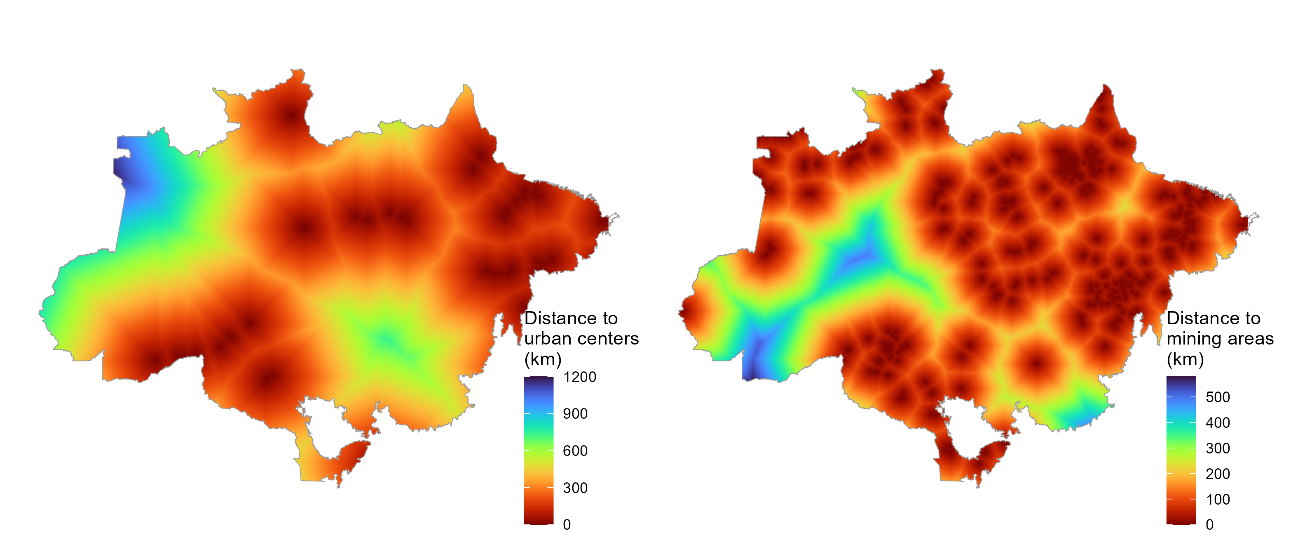

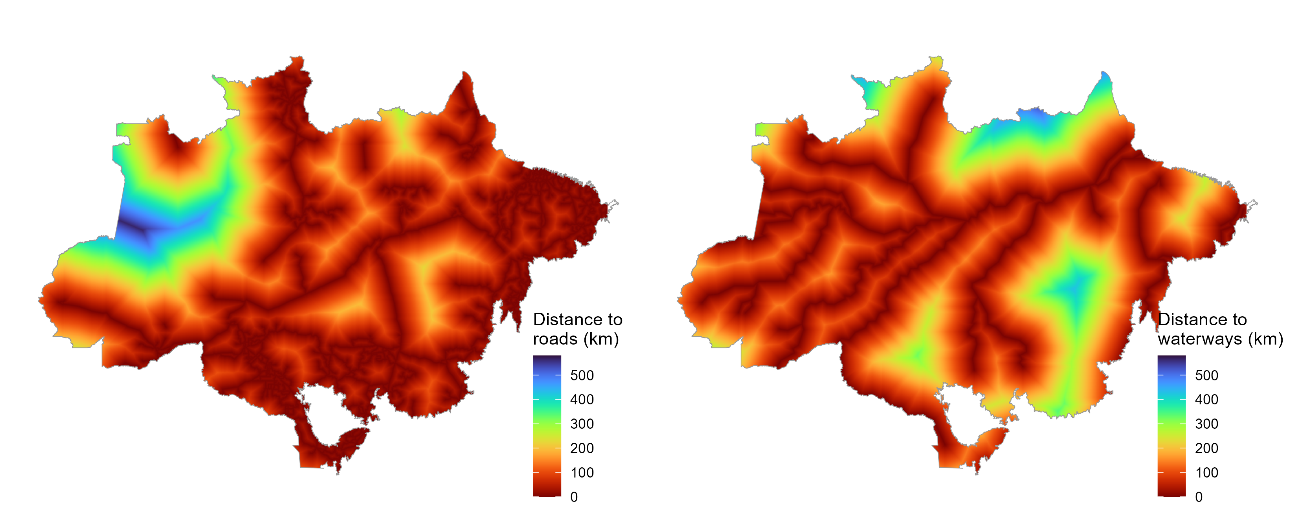

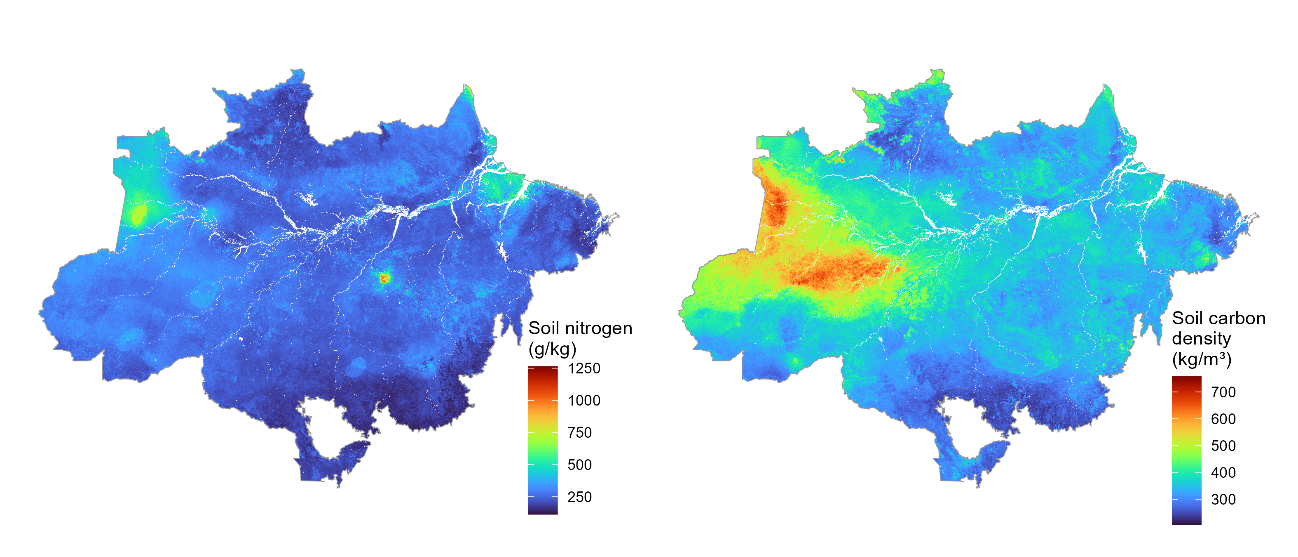

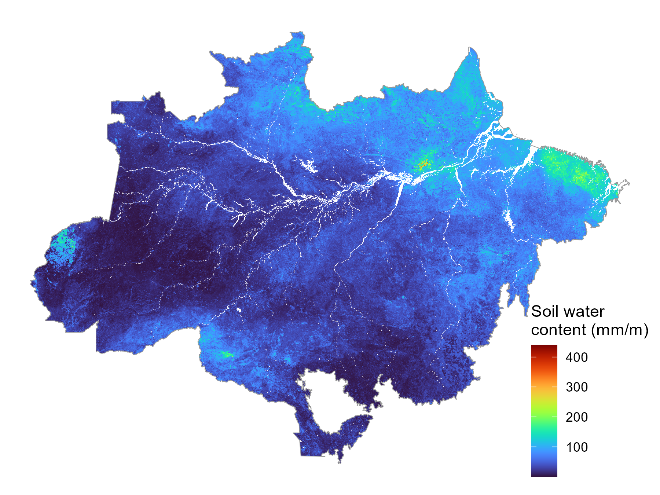

Figure S2. Spatial distribution of the covariates used for statistical matching between protected and unprotected cells within the Brazilian Amazon biome, after data processing and standardization to 1 km resolution (see Table 1 for variable descriptions and sources).

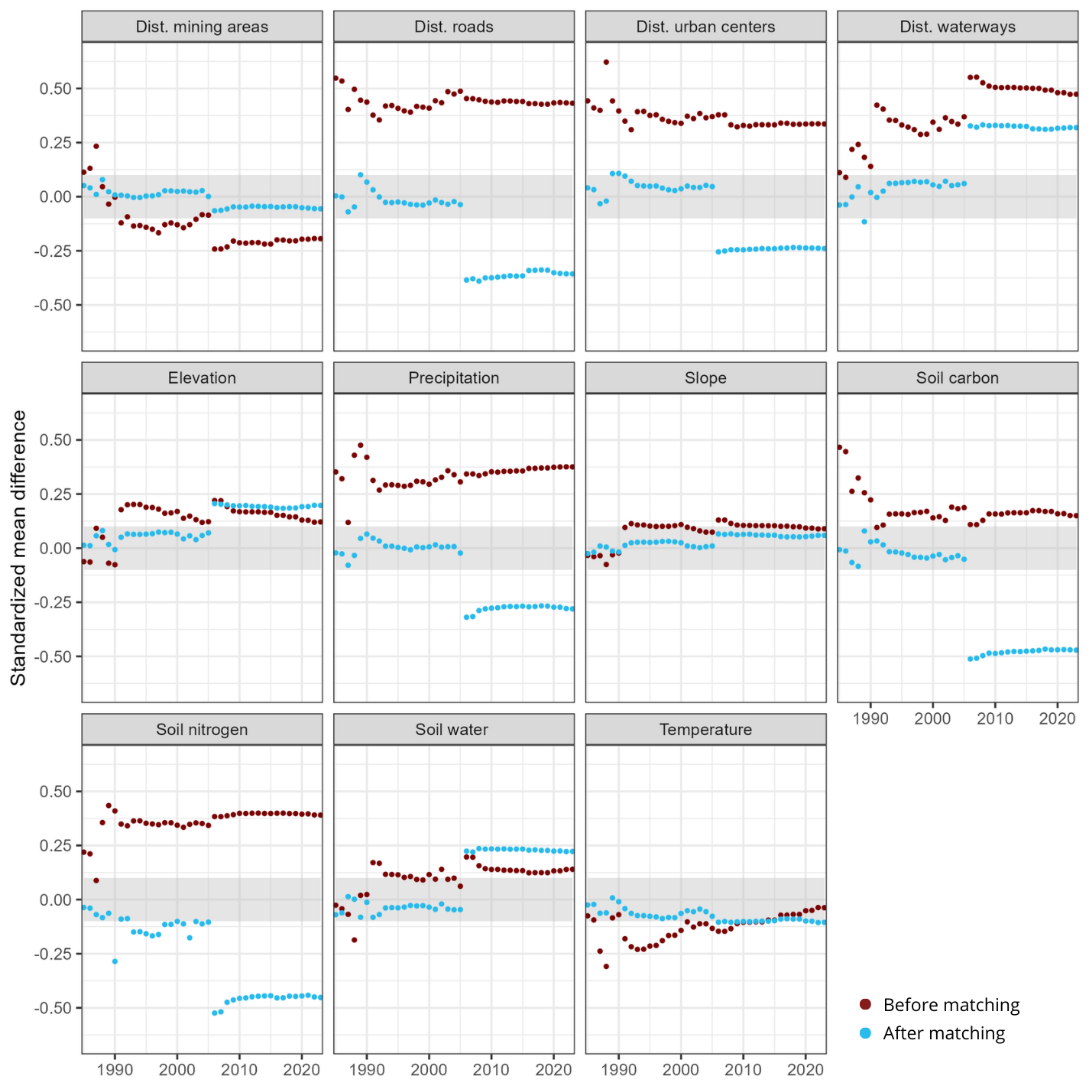

Figure S3. Standardized mean difference between protected areas and control cells, before and after statistical matching, for each of the variables used. The grey zone represents the 0.1 interval. The land use and land cover variable for 1985 was incorporated into the matching procedure through exact matching, ensuring that treated and control units were compared only within the same land cover class; consequently, there is no difference between the values of matched and control cells for this variable, and its corresponding plots were therefore not included.

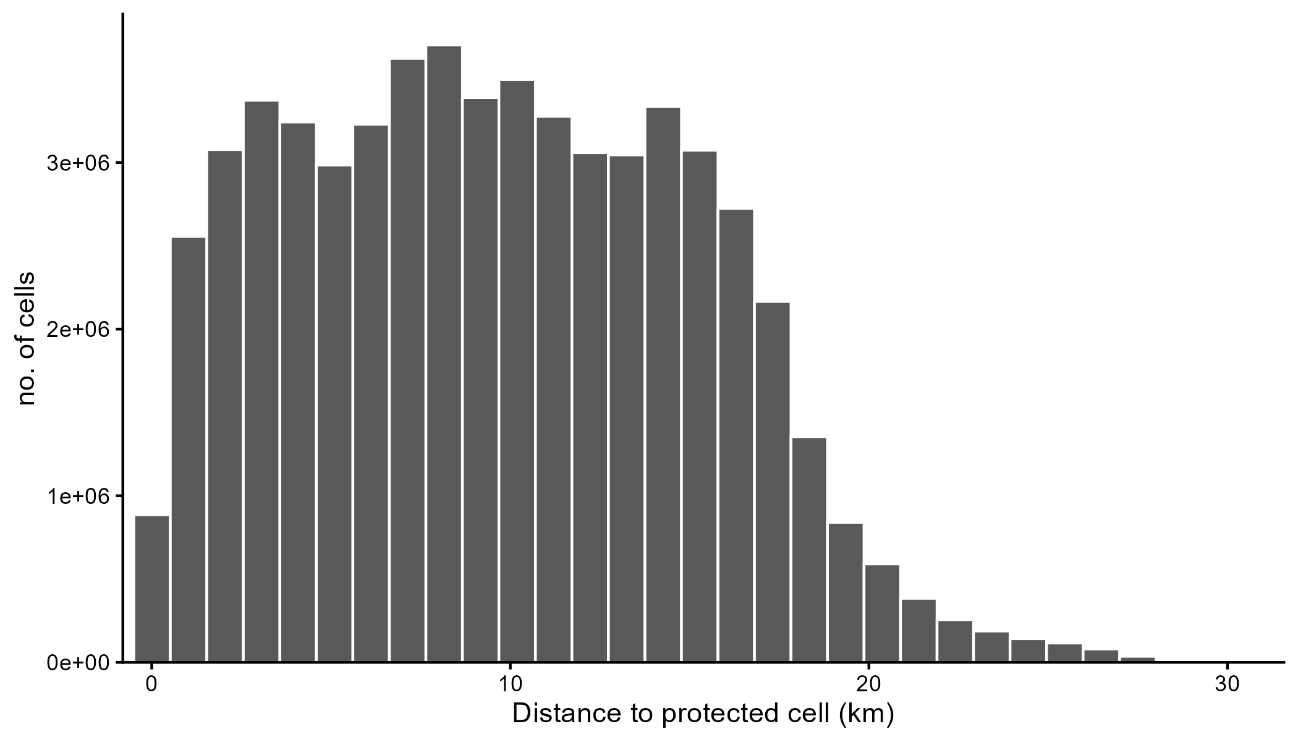

Figure S4. Distribution of Euclidean distances between paired protected and non-protected (control) cells after generalized full matching. Each protected cell was paired with the control cell from the same subclass with the highest estimated probability of receiving protection.

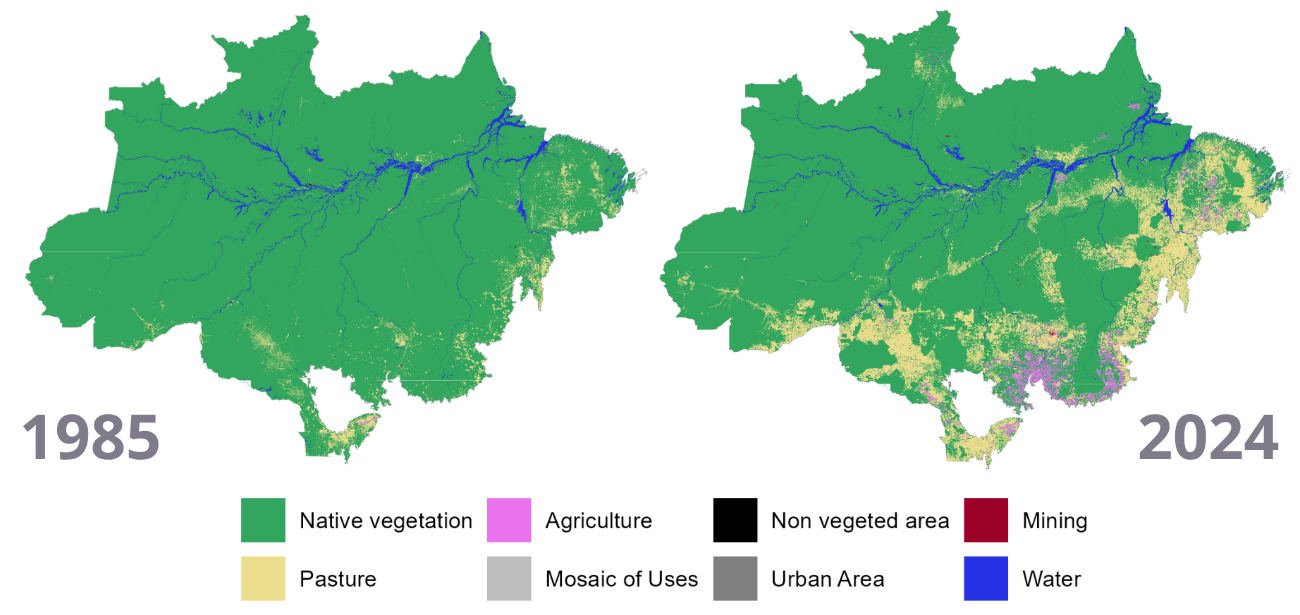

Figure S5. Land use and land cover in the Amazon biome in 1985 (left) and 2024 (right), representing the initial and final states of the analysis period. The maps highlight the expansion of anthropogenic land uses over the period. The original data at 30 x 30 m resolution from the MapBiomas Project (Collection 10) were resampled to a resolution of 1 x 1 km.

Table S2. Distribution of protected areas across impact rate categories by protection type. Impact rate was calculated by dividing total avoided deforestation by area size (km²) and age (years). Categories were defined as: negative impact (rate < 0), low positive impact (rate between 0 and the lower quartile), medium impact (rate between the lower quartile and the median), and high impact (rate above the median)

|  | Indigenous lands | Quilombola Territories | Conservation Units | Total |
| --- | --- | --- | --- | --- |
| No impact | 19 | 49 | 81 | 149 |
| Negative impact | 14 | 21 | 36 | 71 |
| Low positive impact (<.5%) | 91 | 15 | 75 | 181 |
| Medium positive impact (<.8%) | 84 | 18 | 98 | 200 |
| High positive impact (>.8%) | 109 | 18 | 74 | 201 |
| Total | 317 | 121 | 364 | 802 |

Table S3. Regression results for avoided deforestation rate in Brazilian protected areas. Predictors include observed deforestation rate, native vegetation cover, protected area type, mean distance to municipalities, area size (km²), and age of the protected area. State fixed effects are included. Standard errors are heteroskedasticity-consistent (HC1), clustered by state. CU = conservation unit; QT = Quilombola territory; Indigenous land is the reference category and is therefore omitted from the table. State abbreviations refer to Brazilian states: AC = Acre, AP = Amapá, MA = Maranhão, MT = Mato Grosso, PA = Pará, RO = Rondônia, RR = Roraima, TO = Tocantins; Amazonas is the reference category and is therefore omitted from the table.

| **Predictors** | **Estimates** | **Std. error** | **CI** | **p** |
| --- | --- | --- | --- | --- |
| (Intercept) | 0.169 | 0.114 | -0.055 – 0.393 | 0.138 |
| Observed deforestation rate | -0.718 | 0.135 | -0.984 – -0.453 | **<0.001** |
| % of original native vegetation | 0.006 | 0.001 | 0.004 – 0.009 | **<0.001** |
| State [AC] | 0.174 | 0.020 | 0.136 – 0.213 | **<0.001** |
| State [AP] | -0.318 | 0.062 | -0.439 – -0.197 | **<0.001** |
| State [MA] | 0.198 | 0.176 | -0.148 – 0.545 | 0.261 |
| State [MT] | 0.151 | 0.022 | 0.107 – 0.194 | **<0.001** |
| State [PA] | -0.225 | 0.064 | -0.351 – -0.099 | **<0.001** |
| State [RO] | 0.167 | 0.067 | 0.036 – 0.299 | **0.013** |
| State [RR] | -0.182 | 0.039 | -0.258 – -0.106 | **<0.001** |
| State [TO] | -0.155 | 0.166 | -0.481 – 0.170 | 0.349 |
| PA type [QT] | -0.060 | 0.091 | -0.238 – 0.119 | 0.513 |
| PA type [CU] | -0.087 | 0.057 | -0.199 – 0.025 | 0.127 |
| Mean distance to urban centers | -0.0004 | 0.000 | -0.001 – -0.000 | **0.010** |
| Protected are size (km²) | 0.000 | 0.000 | -0.000 – 0.000 | 0.194 |
| Protected are age | -0.002 | 0.003 | -0.007 – 0.003 | 0.447 |
| Observations | 725 | | | |
| R^2^ / R^2^ adjusted | 0.692 / 0.685 | | | |
